# Epigenetic age acceleration is associated with contaminant exposure in common dolphins (*Delphinus delphis*)

**DOI:** 10.64898/2026.06.22.733504

**Authors:** AC Lattmann, EMF Hanninger, EL Betty, X Shen, MJ Anderson, S Gaw, SS Mann, W Gao, KJ Peters, S Yi, JWJ Jokela, KA Stockin

**Affiliations:** New Zealand Institute for Advanced Study, Massey University, Auckland 0745, New Zealand; Department Biology, ETH Zurich, Swiss Federal Institute of Technology, Switzerland; Department of Civil and Environmental Engineering, University of Auckland, Auckland 1142, New Zealand; School of Physical and Chemical Sciences, University of Canterbury, Christchurch, New Zealand; Marine Vertebrate Ecology Lab, Environmental Futures, School of Science, University of Wollongong, Wollongong, Australia; Department of Chemical and Materials Engineering, University of Auckland, Auckland 1142, New Zealand; Department of Environmental Sciences, ETH Zurich, Swiss Federal Institute of Technology, Switzerland

**Keywords:** Biological ageing, PFAS, Heavy metals, Bioaccumulation, Cetaceans

## Abstract

Metals and per- and polyfluoroalkyl substances (PFAS) represent a significant environmental concern, yet their association with epigenetic age acceleration (EAA) remain largely understudied in marine mammals. Here, associations between EAA in common dolphins (*Delphinus delphis*) and life history (sex and sexual maturity), trace metals, and PFAS were investigated. EAA was calculated as the residual in the regression of epigenetic age *vs* chronological age, hence providing a direct measure of the deviation of the epigenetic age of an organism (positive or negative) by comparison with expectation, given their actual chronological age. Sixteen trace elements were quantified in hepatic and renal tissues (*n* = 53). In addition, 28 PFAS were quantified in hepatic tissue (*n* = 58). Associations between EAA and explanatory variables were assessed using regression-based and multivariate modelling approaches (linear models and canonical analysis of principal coordinates). No effect of sex was observed, although sexual maturity did significantly increase EAA. Exposure to metals was significantly associated with EAA, explaining 55.4% of the variation, with hepatic metals (Se, Zn, Cu, Al, Mn) driving this relationship. Although EAA was not significantly related to the total PFAS exposure overall, a subset of PFAS variables (PFBA, PFDA, PFHxS-B, PFNA) showed significant association with EAA after adjusting for sex and sexual maturity. Together, these subsets of metal and PFAS variables, in addition to the selenium-to-mercury (Se:Hg) molar ratio, explained 66.7% of the variation in EAA. Our results identify sexual maturity and specific contaminant mixtures as key potential drivers of EAA in common dolphins, highlighting the possible use of EAA as a biomarker of environmental and physiological stress in marine mammals.

## Introduction

Marine environments are often the sink for anthropogenic contamination, with pollutant levels in these environments projected to increase further due to ongoing population growth, the temporal persistence of pollutants, and increasing global production (Shahidul Islam & Tanaka, 2004; Willis et al., 2022). Among the various pollutants in the marine environment, metals represent one of the greatest toxicological threats due to their persistence, capacity for bioaccumulation, and potential for biomagnification (Gao & Chen, 2012; Jiménez-Ballesta et al., 2018; Sujitha et al., 2019; Zhou et al., 2007). Additionally, the widespread contamination of groundwater and surface water by per- and polyfluoroalkyl substances (PFAS) is a major environmental concern (Evich et al., 2022; Roth et al., 2021).

Due to their longevity and position at the top of marine food webs, cetaceans (whales, dolphins and porpoises) are especially susceptible to the bioaccumulation and biomagnification of metals through dietary intake (Boon et al., 1994; Garcia-Cegarra et al., 2020). Consequently, metals such as mercury (Hg) and cadmium (Cd) can reach elevated concentrations in these species (Caurant et al., 1996; Hansen & Danscher, 1995; López-Berenguer et al., 2020), rendering cetaceans valuable bioindicators of marine contamination (Cáceres-Saez et al., 2018; Dehn et al., 2006; Mapunda et al., 2017; Seixas et al., 2007, 2009; Xiong et al., 2019). Similarly, reports have documented the bioaccumulation and biomagnification of PFAS in top predators such as cetaceans (Androulakakis et al., 2022; Fair et al., 2013), further supporting their use as sentinels of environmental contamination (Alava et al., 2020; Bossart, 2011; Ross, 2000).

In this context, assessing how contaminant exposure influences biological ageing in cetaceans requires metrics that capture inter-individual variation in ageing rates, such as epigenetic age acceleration (EAA). Epigenetic clocks, based on DNA methylation patterns, provide a powerful method for estimating biological or “epigenetic age” (Felix & Cecil, 2019; Shanthikumar et al., 2020). Based on epigenetic clocks, EAA can then be derived as residuals from regressing DNAm-predicted age (DNA methylation age) on chronological age (Nwanaji-Enwerem et al., 2018), reflecting the rate of epigenetic ageing relative to chronological age (Jylhävä et al., 2017). A positive EAA indicates that an individual is ageing faster biologically than expected for their chronological age, whereas a negative EAA indicates slower biological ageing compared to chronological age (Horvath & Raj, 2018).

In humans, several studies have linked differences in biological age to variation in trace metals (Ryoo et al., 2025; Wang et al., 2025). Specifically, a mixture of nonessential metals (Tungsten (W), Arsenic (As), Cadmium (Cd)) was associated with EAA, whereas a subset of essential metals (Selenium (Se), Zinc (Zn), Molybdenum (Mo)) reportedly lowered EAA (Boyer et al., 2023). Cadmium, in particular, was associated with increased EAA across multiple published methylation-based epigenetic clocks, including those that capture mortality risk (GrimAge), biological ageing as well as health risks (PhenoAge) and the pace of ageing (DunedinPACE) (Boyer et al., 2023). Evidence indicates that trace metal exposure can alter DNA methylation patterns (Elkin et al., 2022; Kim et al., 2024), suggesting a plausible mechanistic link between metal exposure and epigenetic ageing; however, direct empirical studies quantifying EAA in wildlife exposed to trace metals remain limited.

Associations between PFAS exposure and biological ageing in humans remain inconsistent, with reports of age acceleration (Khodasevich et al., 2025; Yan et al., 2025; Zhao et al., 2024), age deceleration (Chaney & Wiley, 2023) or no association between PFAS exposure and EAA (Xu et al., 2020). For instance, a study of 59 Swedish women exposed to PFAS-contaminated drinking water identified differential methylation at 117 cytosine–phosphate–guanine (CpG) sites but found no association with EAA (Xu et al., 2020). In contrast, Khodasevich et al. (2025) reported positive correlations between PFAS exposure, including individual compounds such as Perfluorononanoic acid (PFNA) and Perfluorooctanesulfonic acid (PFOS), as well as the total PFAS mixture, and EAA in males. Complementing these findings in humans, a study of 197 firefighters (89.3% male; mean age 38.6 years) reported that exposure to specific PFAS, including perfluorohexane sulfonate (PFHxS), perfluorooctanoate (PFOA), and the sum of branched perfluorooctane sulfonate isomers (Sm-PFOS), was associated with EAA (Goodrich et al., 2021). While most studies have focused on humans, the effects of trace metal and PFAS exposure on EAA in marine mammals remain poorly understood. To date, no studies have directly examined associations between trace metals and EAA and only one study (Peters et al., 2026) has examined total PFAS concentrations in the context of epigenetic ageing in cetaceans. Accordingly, current understanding of metals, PFAS and EAA relationships are extremely limited and somewhat preliminary. Moreover, the combined effects of trace metals and PFAS on EAA in the context of sex and sexual maturity are yet to be investigated in marine mammals.

We examined EAA in relation to contaminant exposure in common dolphins (*Delphinus delphis*). Life history parameters, including sex and sexual maturity, were included to assess their potential influence on EAA and to account for their effects in a multivariate model. We applied ordinary least squares (OLS) regression to evaluate whether EAA varies by sex or with sexual maturity in common dolphins. Multiple regression models (achieved *via* canonical analysis of principal coordinates and distance based linear models) were used to test the association of EAA *vs* (1) hepatic and renal trace metal concentrations; and (2) accumulated hepatic PFAS. Specifically, we identified which metals and PFAS most strongly influenced EAA, and incorporated these into a multivariate model together with the selenium-to-mercury (Se:Hg) ratio (indicator of mercury detoxification capacity, Law et al., 2003) and life history parameters (sex and sexual maturity), to investigate their combined influence on EAA.

## Methods

### Sample collection

In total, 75 common dolphins were sampled post-mortem between 2000 and 2023 from bycatch (*n* = 8), single (*n* = 35) and mass (*n* = 32) stranding events. For all individuals, chronological age was estimated from dental growth layer groups (GLGs; Palmer et al., 2023), alongside epigenetic age and EAA. Trace metals (mg/kg) and PFAS concentrations (ng/g) were measured in liver and kidney tissues from two overlapping subsets of individuals (*n* = 53 and *n* = 58, respectively).

### Epigenetic age determination

Epigenetic age estimates were obtained from the species-specific epigenetic clock developed for common dolphins by Hanninger et al. (2025). This clock was calibrated using age estimates derived from dentinal GLGs in tooth sections (Palmer et al., 2023). In brief, DNA methylation profiles from skin samples were generated using the HorvathMammalMethylChip40 array, and epigenetic clocks were constructed and validated using the MammalMethylClock R package (Zoller & Horvath, 2024) with Leave-One-Out Cross-Validation (LOOCV). Additional methodological details are provided in Hanninger et al. (2025).

### Epigenetic age acceleration calculation

EAA (AgeAccelLOO) was calculated as the residual from a linear regression of DNAmAgeLOO (predicted epigenetic age) on chronological (dental) age: *AgeAccelLOO = DNAmAgeLOO − (β₀ + β₁ × Age)*, where β₀ and β₁ denote the estimated intercept and slope (MammalMethylClock R package, Zoller & Horvath, 2024). DNAmAgeLOO systematically overestimated age in younger individuals and underestimated it in older individuals, resulting in a non-zero intercept and slope < 1 (Hanninger et al., 2025).

### Life history

Sex and sexual maturity were assessed *a priori* via gross and histological examination of reproductive organs, as reported by Palmer et al. (2022) and Palmer et al. (2023) for males and females, respectively. In males, sexual maturity was defined by the presence of spermatozoa within the seminiferous tubules and/or epididymides, following Palmer et al. (2022). In females, sexual maturity was defined by the presence of at least one *corpus albicans* (CA) or *corpus luteum* (CL) on an ovary, or by evidence of pregnancy or lactation. In cases where gross morphological or histological assessments were unavailable (*n* = 4), sexual maturity was determined using Gompertz growth curves based on total body length (Palmer et al., 2022, 2023), with classifications subsequently cross-validated through radiographic assessment of pectoral flipper bone development following Hanninger et al. (2026). For two individuals (KS14-56Dd, WS02-40Dd), sexual maturity could not be determined by gonadal examination nor inferred from total body length or pectoral flipper development and were therefore excluded from subsequent analyses.

### Trace element and PFAS contaminant analysis

Liver and kidney samples (∼1 g wet weight) were aseptically collected from 53 common dolphins during post-mortem examinations. The tissue samples were subsequently freeze-dried, and analysed for 16 trace elements following Lischka et al. (2021). Samples underwent microwave-assisted acid digestion prior to inductively coupled plasma mass spectrometry (ICP-MS; Agilent 8900) analysis. Mercury preparation followed Aldridge et al. (2017). Quality assurance included analytical blanks, fish protein certified reference material (DORM-4), and duplicate analysis of 10% of samples. Element concentrations were calculated on a dry weight basis and converted to wet weight concentrations using organ-specific moisture conversion factors (See Supplementary Materials for full methods description).

A total of 58 dolphin liver samples were analysed for 28 PFAS variables (see Supplementary Materials, Table S1, for the full list of analytes and their CAS numbers) using a modified extraction and liquid chromatography–tandem mass spectrometry (LC–MS/MS) method following Shen et al. (2026) and U.S. EPA (2024). Briefly, approximately 1 g of freeze-dried liver tissue was homogenised, spiked with isotope-labelled internal standards prior to extraction, and then subjected to a three-step solvent extraction using methanol (including KOH) and acetonitrile. Extracts were combined, concentrated under a stream of nitrogen, and purified using ENVI-Carb and weak anion exchange (WAX) solid-phase extraction (SPE) cartridges. Quantification was performed using an Agilent 1290 Infinity II LC system coupled to a 6460C triple quadrupole mass spectrometer operating in negative electrospray ionisation mode with multiple reaction monitoring (MRM). The analytical method targeted perfluoroalkyl carboxylic acids (C4–C12), perfluoroalkane sulfonic acids (C4–C8), ether PFAS, and selected fluorotelomers. External calibration standards ranged from 0 to 7.5 ng/mL, and the limit of quantification (LOQ) was 0.2 ng/mL, corresponding to a limit of reporting (LOR) of 0.2 ng/g wet weight in tissue (see Supplementary Materials for a more complete description of these methods. MRM transitions for the 28 target PFAS analytes are provided in Supplementary Materials, Table S2).

### Statistical analyses

#### Epigenetic age acceleration (EAA)

EAA was calculated using the epigenetic clock calibrated on the full dataset of 75 individuals (Hanninger et al., 2025). Although Hanninger et al. (2025) developed multiple clocks using different subsets of the data (*relaxed, strict, restricted*), the full-sample clock (*relaxed*) was selected for the present study because it captured the greatest epigenetic age variation across the dataset as a whole, including individuals exhibiting both accelerated and decelerated epigenetic ageing. Details of the *strict* and *restricted* subsets are presented in the Supplementary Materials (Fig S1).

#### EAA vs life history

Ordinary least squares (OLS) regression was done (using R version 4.4.1, R Core Team, 2026) of EAA *vs* sex and sexual maturity. This analysis was repeated, changing the order of the terms, to identify overlap in the amount of variation in EAA that could be explained by these two variables.

#### Influence of metals on EAA

##### Pre-treatment of metal variables

Values below the limit of detection (LOD) for metal concentrations were imputed as LOD/√2, following Croghan & Egeghy (2003). Imputation was performed separately for each combination of metal, organ, and analytical batch to account for variation in LODs across these factors (LOD values are provided in Supplementary Tables, Table S3 and Table S4). Metal concentration variables were strongly right-skewed and were therefore log-transformed prior to analysis. Only samples with complete data from both liver and kidney were included (*n* = 53). Threshold-metric multidimensional scaling (tmMDS) ordination of a Euclidean distance matrix based on the normalized log-transformed metal variables showed a pronounced analytical batch effect (Supplementary Materials, Fig S2). Thus, for ensuing multivariate analyses using canonical analysis of principal coordinates (CAP), a residualised distance matrix that removed batch effects was obtained using the PERMANOVA routine (Anderson, 2001a, 2017), implemented in PRIMER 8 with PERMANOVA+ (Anderson & Euinton 2026). In addition, univariate residuals were obtained by fitting a one-way ANOVA *vs* ‘batch’ as a linear model (lm) in R (R Core Team, 2026) to each normalized log-transformed metal variable. These batch-adjusted metal values were then used as predictor variables in subsequent multiple regression models (CAP and DISTLM).

##### Multivariate association between trace metals and EAA

To evaluate the relationship between EAA and overall metal concentrations across hepatic and renal tissues (*p* = 32 variables, 16 metals measured per tissue), a canonical analysis of principal coordinates (CAP; Anderson & Robinson, 2003; Anderson & Willis, 2003) was done using the CAP routine in PRIMER 8 with PERMANOVA+ (Anderson & Euinton, 2026). Analyses were based on a Euclidean distance matrix calculated from batch-adjusted, normalized log-transformed metal concentrations. Here, CAP was used to find an axis (CAP1), which is a linear combination of principal coordinate (PCO; Gower, 1966) axes derived from the metals distance matrix, so as to maximise the correlation between EAA and CAP1. The canonical correlation (δ1) indicates the proportion of the variance in EAA explained by the CAP axis. To avoid overfitting, only a subset of PCO axes was included. The optimal number of PCO axes (*m*) was determined by minimising the leave-one-out residual sum of squares of the CAP model. For trace metals and EAA, m = 9 was identified as optimal. A plot of EAA vs CAP1 was generated to visualise the relationship. Statistical significance of the CAP model was assessed using a permutation test for m = 9 with 9,999 permutations.

##### Trace metals most strongly associated with EAA

The CAP analysis showed clear redundancy across the metal variables in their ability to explain variation in EAA (32 metal variables were reduced to m = 9 PCO axes in the CAP model), due to their inter-correlations. It was desirable to identify a useful subset of metal variables for subsequent overall models of EAA that would include other factors as well (i.e., sex, maturity and PFAS information). Distance-based linear modelling (DISTLM; McArdle & Anderson, 2001) was used to implement multiple regression to identify key metal predictors (out of 32 variables consisting of the batch-adjusted, normalised, log-transformed liver and kidney metal concentrations) of EAA. A DISTLM on a Euclidean distance matrix for one variable (here, EAA) is equivalent to performing a classical multiple regression (for the fit), but with all tests done using appropriate permutation algorithms. Marginal tests of the relationships between each batch-adjusted metal variable and EAA were done using 9,999 permutations as an initial exploratory step.

Next, a step-wise selection procedure on the basis of the small sample-size corrected Akaike information criterion (AICc; Burnham & Anderson, 2002) was used to identify a parsimonious subset of metal variables. (Finding an AICc ‘best’ model was infeasible to achieve with 32 variables, corresponding to over 4.29 billion possible models). Sequential tests of individual variables in the resulting step-wise AICc subset were then done using 9999 permutations under a reduced model (Anderson, 2001b; Freedman & Lane, 1983). Pearson correlations were calculated between EAA and individual variables identified by the step-wise AICc analysis to clarify the nature of these relationships.

#### EAA vs PFAS concentrations

A total of 28 PFAS variables were measured. As PFAS concentrations were right-skewed, values were log-transformed using [inline] log(x + LOQ/√2) prior to analysis, with the LOR at 0.2 ng/g for all PFAS components. PFAS data were filtered to include only individuals with detectable measurements (*n* = 58).

To investigate the relationship between EAA and PFAS variables, CAP and DISTLM analyses were done in the same manner as described above for the metal variables. The CAP analysis was based on *m* = 10 principal coordinate axes (selected using leave-one-out residual sum of squares) derived from a Euclidean distance matrix of 28 PFAS variables, enabling evaluation of the overall relationship between PFAS profiles and EAA. DISTLM (both marginal tests and model selection based on the AICc criterion) was used to explore the relationships between EAA and individual PFAS variables (*n* = 28), and to obtain a parsimonious subset of PFAS variables to include in subsequent overall models of EAA.

#### Combined analysis of EAA vs factors and predictor variables

A CAP analysis was performed to visualize multivariate associations between multiple predictors (metal subset selected via DISTLM, PFAS subset selected via DISTLM, Se:Hg ratio) and EAA. The effect of sexual maturity on EAA was removed using PERMANOVA, and the resulting residuals were used in the CAP analysis. Nine PCO axes (*m* = 9) were selected, as this number minimised the leave-one-out residual sum of squares of the CAP model (Anderson, 2008). A CAP ordination plot was then used to visualise the association between overall PFAS concentrations and EAA. Statistical significance of the CAP model was assessed using a permutation test with 9,999 permutations.

Metal and PFAS subsets showing the strongest associations with EAA, identified using DISTLM (stepwise selection based on AICc), were incorporated into a combined model to assess their joint influence on EAA (based on R²). The model additionally included sex, sexual maturity, and the selenium-to-mercury (Se:Hg) molar ratio as covariates, with sex and sexual maturity coded as binary variables (−1/1 coding). The dataset (n = 51) was restricted to individuals with complete data across all variables. Variables were entered in a specified order, with biological predictors (sex and sexual maturity) included first, followed by the Se:Hg molar ratio, the selected metal subset, and the PFAS subset. This hierarchical structure allowed partitioning of variation in EAA attributable to contaminant exposure (metals and PFAS) and Se:Hg, while controlling for variation associated with sex and sexual maturity.

## Results

### Influence of life history on epigenetic age acceleration

The sample was composed of 44% (*n* = 33) and 56% (*n* = 42) males and females, respectively. Of the males, 45.5% (*n* = 15) were sexually immature, while 33.3% (*n* =14) of females were sexually immature. After adjusting for sexual maturity, males showed a non-significant increase in EAA compared to females (β = 1.04, 95% CI [−0.50, 2.58], *p* = 0.18, HC1 robust standard errors). After adjusting for sex, sexually mature individuals exhibited significantly higher EAA compared to immature individuals (β = 2.24, 95% CI [0.92, 3.56], *p* = 0.001, HC1 robust standard errors; Fig 1).

**Fig 1.**
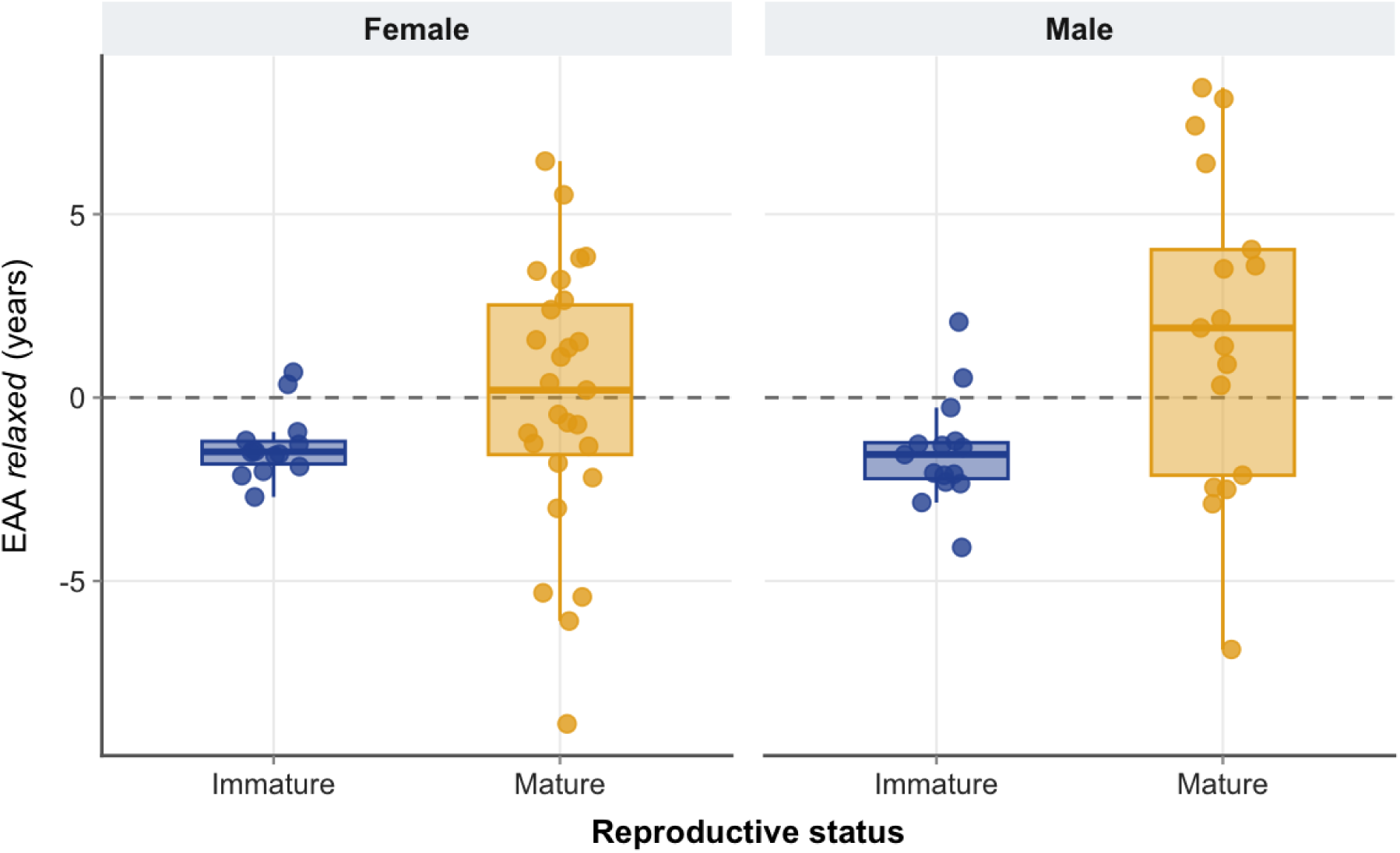
Epigenetic age acceleration (EAA *relaxed*; years) in New Zealand common dolphins (*Delphinus delphis*) by sex and reproductive status. Panels are stratified by sex (male and female), with sexual maturity indicated within each sex. Boxplots show the median and interquartile range of EAA *relaxed*, with individual observations overlaid as jittered points. Colours indicate reproductive status: immature (blue) and mature (orange).

### Influence of trace metals on EAA

Collectively, 71% (*n* = 53) of the dolphins had measured burdens of 16 metals across both liver and kidney. Detection frequency across both tissues ranged from 6.9% (Pb) to 100% (Mg, Cu, As, Se, and Hg), with mean concentrations among detected samples ranging from 0.06 ± 0.06 mg/kg (V) to 143.2 ± 36.4 mg/kg (Mg). A CAP analysis testing combined effects of metal concentrations in liver and kidney on EAA was statistically significant (*m* = 9), with a canonical correlation of δ = 0.554 and *p* = 0.048 (based on 9,999 permutations; Fig 2).

**Fig 2.**
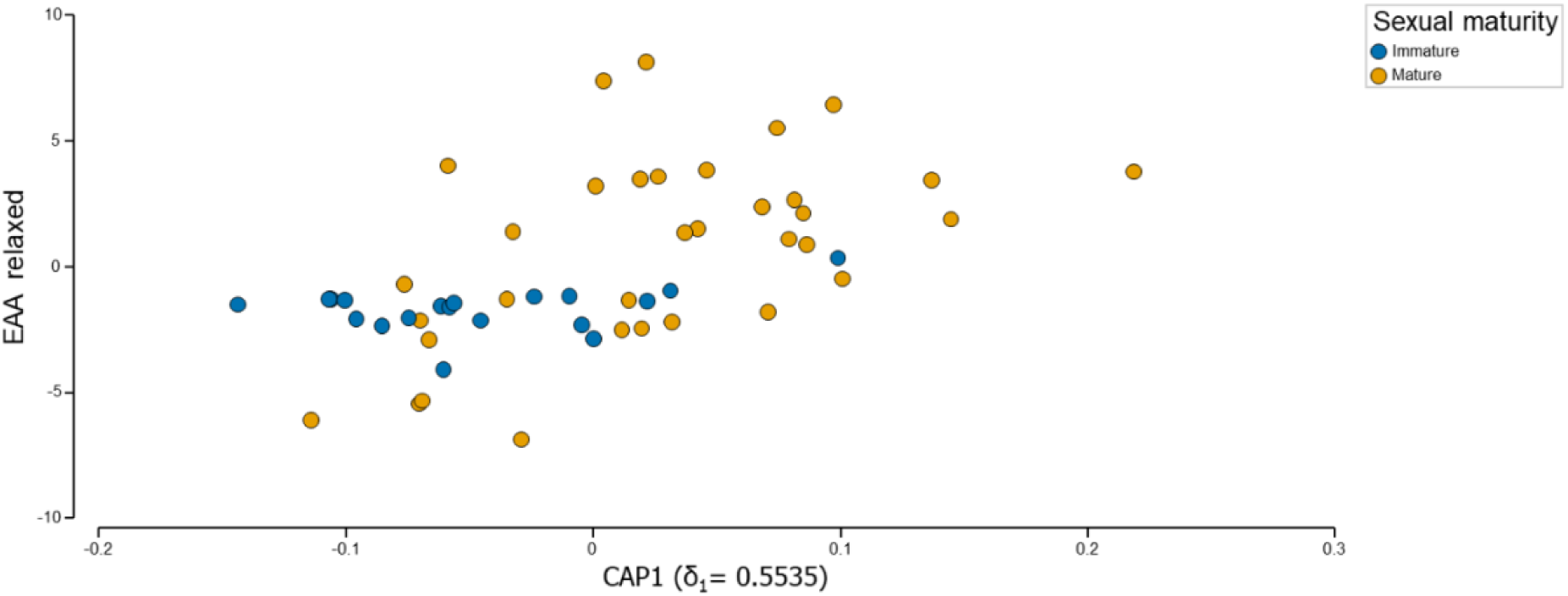
CAP analysis based on *m* = 9 PCO axes derived from a Euclidean distance matrix, built from normalised, log-transformed, batch-adjusted, metal concentrations measured in liver and kidney (p = 32 variables; 16 per organ) of New Zealand common dolphins (*Delphinus delphis*, *n* = 53 individuals), showing their multivariate association with epigenetic age acceleration (EAA). CAP1 represents the first canonical axis, and δ1 denotes the canonical correlation between CAP1 and EAA. Individual dolphins are coloured according to sexual maturity status, with sexually immature individuals shown in blue and sexually mature individuals shown in orange.

#### Trace metals most strongly associated with EAA

Individually, selenium in liver (Se.L) explained the largest proportion of variation in EAA (12.77%, *p* = 0.007), followed by cadmium in liver (Cd.L; 11.19%, *p* = 0.014), mercury in liver (Hg.L; 10.76%, *p* = 0.017), and mercury in kidney (Hg.K; 7.37%, *p* = 0.048), all of which were statistically significant. Cadmium in kidney (Cd.K; 7.26%, *p* = 0.055) and magnesium in kidney (Mg.K; 6.47%, *p* = 0.064) approached significance and still displayed relatively high contributions to variation (Supplementary Materials, Table S5).

Stepwise model selection using AICc yielded a parsimonious subset of five liver metals: selenium (Se.L), zinc (Zn.L), copper (Cu.L), aluminium (Al.L), and manganese (Mn.L), which together explained 35.98% of the variation in EAA. Sequential tests showed Se.L as the primary contributor (12.77%, *p* = 0.009), followed by Zn.L (20.44%, +7.67%, *p* = 0.032), Cu.L (27.90%, +7.46%, *p* = 0.029), Al.L (32.57%, +4.67%, *p* = 0.075), and Mn.L (35.98%, +3.41%, *p* = 0.121) (Supplementary Materials, Table S6).

#### Direction of association of liver trace metals with EAA

Metals displayed variable-specific direct associations between hepatic contaminant burden and EAA. Zinc (Zn.L; *r* = 0.181) and selenium (Se.L; *r* = 0.357) were positively associated with EAA, indicating that higher concentrations of these elements were associated with increased EAA. In contrast, aluminium (Al.L; *r* = −0.155), manganese (Mn.L; *r* = −0.046), and copper (Cu.L; *r* = −0.132) showed negative direct associations with EAA, suggesting that higher concentrations of these metals were associated with reduced EAA. Overall, selenium exhibited the strongest positive association with EAA among the metals examined.

### Influence of PFAS on EAA

PFAS (*n* = 28) were measured across 58 individuals, including 31 females and 27 males. Detection frequency across compounds ranged from 0 to 84.5% (PFOS-N), with five compounds (PFPeA, 4:2FTS, HFPO-DA-CO2, NFDHA-CO2, and NFDHA) remaining below the LOR (0.2 ng/g for all samples). Mean concentrations among detected compounds ranged from 0.21 ± 0.01 ng/g (PFEESA) to 66.3 ± 109.9 ng/g (PFOS-N).

#### Multivariate association between PFAS and EAA

Overall levels of PFAS (total of 28 compounds) was not significantly associated with EAA (CAP, m = 10; δ = 0.522; *p* = 0.107 (Supplementary Materials, Fig S3). However, in marginal tests, PFBA explained significant variation in EAA (12.33%, *p* = 0.032) and was the strongest single predictor. PFOA-N was also statistically significant, explaining 7.33% of the variation (*p* = 0.039). Several other compounds, including PFHxA (5.56%, *p* = 0.080), PFDA (5.25%, *p* = 0.082) and PFHxS-B (5.12%, *p* = 0.094), had moderate associations, but these were not statistically significant (*p* > 0.05). All remaining PFAS explained less than 1.6% of the variation and were non-significant (*p* > 0.10) (Supplementary Materials, Table S7).

The step-wise AICc model included four PFAS variables: PFBA, PFDA, PFHxS-B, and PFNA, which together explained 27.84% of the total variation in EAA. PFOA-N was significant in marginal tests but was not retained in the optimal stepwise AICc model. In the sequential tests, PFBA was chosen first as the most important predictor (12.33%, *p* = 0.037), followed by PFDA, which increased the explained variation to 19.85% (+7.52%, *p* = 0.026). Subsequent additions of PFHxS-B (+4.03%, *p* = 0.094) raised the cumulative explained variation to 23.88%. followed by the addition of PFNA (+3.96%, *p* = 0.092), resulting in a total of 27.84%. Note that the latter two sequential tests had weak evidence against the null hypothesis (0.05 < *p* < 0.10) (Supplementary Materials, Table S8).

Overall, these analyses indicate that PFBA is the primary PFAS compound associated with variation in EAA, with weaker contributions from PFDA, PFHxS-B, and PFNA.

#### Direction of association of PFAS with EAA

Pearson correlation analysis between EAA and selected PFAS indicated variable-specific associations. PFBA and PFHxS-B were excluded from Pearson correlation analysis due to low detection frequencies (1.7% and 3.4%, respectively). PFNA (*r* = 0.074, detection frequency 55.2%) and PFDA (*r* = 0.229, detection frequency 50.0%) both showed positive correlations with EAA.

### Combined analysis of EAA vs factors and predictor variables

#### Multivariate association between hepatic metals, PFAS, Se:Hg ratio, and EAA

A canonical analysis of principal coordinates (CAP) testing the association between EAA and combined liver metal concentrations (Se, Zn, Cu, Al, Mn; subsets identified in DISTLM) and PFAS concentrations (PFBA, PFDA, PFHxS-B, and PFNA; subsets identified in DISTLM), the selenium-to-mercury (Se:Hg) ratio was statistically significant (m = 9; δ = 0.667; *p* = 0.002; 9,999 permutations; Fig 3), indicating a strong multivariate correlation between the predictor variables and EAA. Residuals of sexual maturity were used to account for its effects in the analysis.

**Fig 3.**
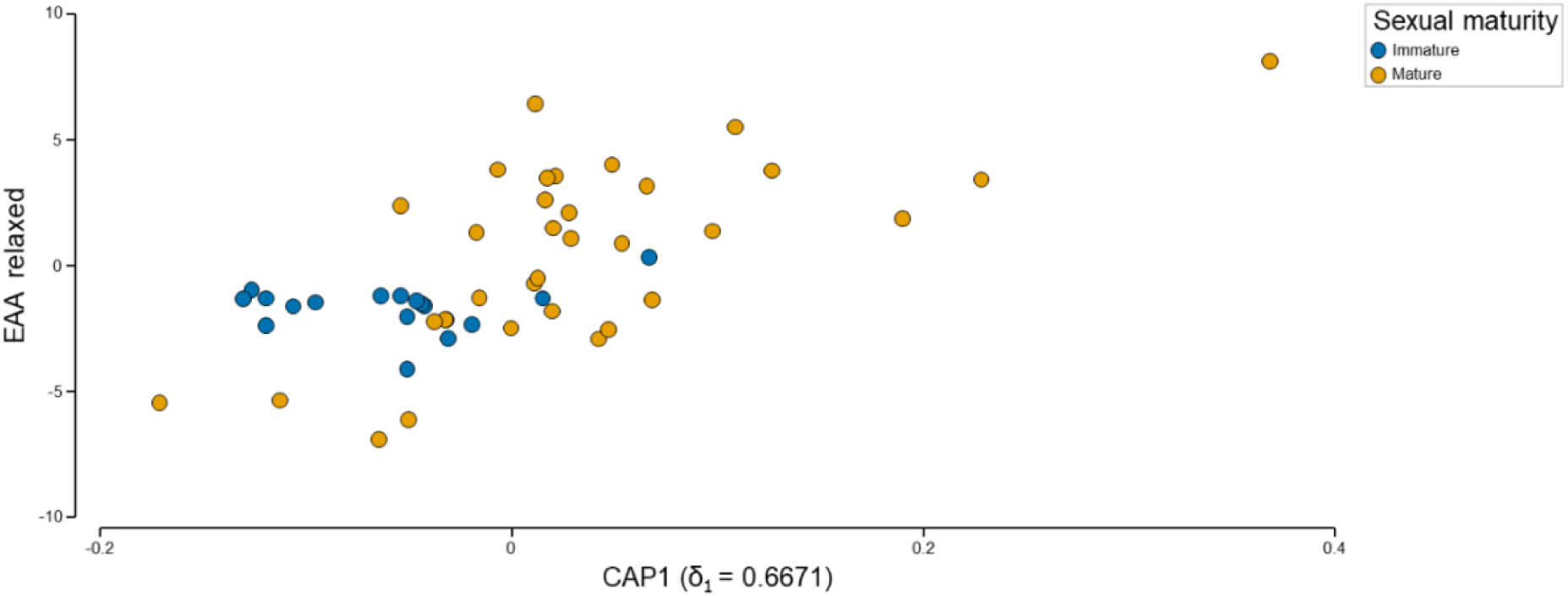
CAP analysis based on *m* = 9 PCO axes derived from a Euclidean distance matrix, built from multiple predictor variables (metal subset, PFAS subset, and Se:Hg ratio) of New Zealand common dolphins (*Delphinus delphis*, *n* = 51), showing their multivariate association with epigenetic age acceleration (EAA). The metal subset included normalised, log-transformed, batch-adjusted concentrations of selenium (Se), zinc (Zn), copper (Cu), aluminium (Al), and manganese (Mn), all measured in liver tissue. The PFAS subset included log-transformed concentrations of PFBA, PFDA, PFHxS-B, and PFNA. Prior to CAP analysis, the effect of sexual maturity was removed using PERMANOVA in PRIMER, and the resulting residuals were used in the CAP analysis. CAP1 represents the first canonical axis, and δ1 denotes the canonical correlation between CAP1 and EAA. Individuals are coloured according to sexual maturity status, with sexually immature individuals shown in blue and sexually mature individuals shown in orange.

#### Metal and PFAS associations with EAA after adjustment for life history variables

Sexual maturity influenced EAA (11.81%, *p* = 0.015), whereas sex alone was not significant (0.76%, *p* = 0.549). While Se:Hg ratio explained 10.24% of the variation in EAA (*p* = 0.080) and an additional 6.73% after adjusting for sex and sexual maturity (*p* = 0.173), neither effect was significant. In contrast, the selected metal and PFAS subsets each individually explained significant portions of the variation in EAA (34.26%, *p* = 0.002 and 27.61%, *p* = 0.004, respectively) (Supplementary Materials, Table S9). Notably, both the metal subset (*p* = 0.021) and PFAS subset (*p* = 0.013) remained statistically significant even after adjusting for the potential effects of sex, sexual maturity and the Se:Hg ratio. All five variable groups together explained 58.08% of the total variation in EAA (*R*² = 0.581) (Table 1).

**Table 1.** Sequential tests of final overall model (Specified order, *R*²) Results of distance-based linear models (DistLM) showing the sequential contribution of sex, sexual maturity, a liver metal subset (Se, Zn, Cu, Al, Mn), a PFAS subset (PFBA, PFDA, PFHxS-B, PFNA), and the selenium-to-mercury (Se:Hg) molar ratio in explaining variation in epigenetic age acceleration (EAA) in New Zealand common dolphins (*Delphinus delphis*). The analysis was conducted using a forward selection procedure based on *R*², with predictors entered in a specified order. Values represent the additional proportion of variation explained at each step (%) and the cumulative *R* ². Associated P-values are provided for each step. Statistically significant predictors (p < 0.05) are highlighted in bold font.

| Variable | P-value | % Added | Cumulative (%) |
| --- | --- | --- | --- |
| Sex | 0.542 | 0.76 | 0.76 |
| **Sexual maturity** | **0.018** | **11.17** | **11.93** |
| Se:Hg ratio | 0.173 | 6.73 | 18.66 |
| **Metals subset** | **0.021** | **21.91** | **40.57** |
| **PFAS subset** | **0.013** | **17.51** | **58.08** |

## Discussion

We provide here the first evidence that trace metal exposure influences EAA in a cetacean species. We also explore, for the first time, the combined effects of trace metals and PFAS on EAA in any cetacean species.

When the contaminant subsets were combined with the selenium-to-mercury (Se:Hg) ratio, the overall model showed a strong multivariate association with EAA (*R*^2^ = 58.08%), indicating that combined exposure variables explain substantially more variation in EAA than trace metals (*R* ^2^ = 35.98%) or PFAS (*R* ^2^ = 27.84%) alone. Meanwhile, while no effect of sex on EAA was evident, sexually mature individuals did exhibit, on average, higher EAA compared with immature conspecifics.

The sixteen hepatic and renal trace elements, when considered as a group, showed a significant association with EAA in common dolphins, explaining 55.4% of the variation. A subset of hepatic metals (Se, Zn, Cu, Al, Mn) showed the strongest associations, explaining 35.98% of the variation, suggesting that only specific metals primarily drive variation in EAA. Notably, the strongest associations in the present study were observed for metals measured in liver tissue. This suggests that hepatic accumulation may be particularly relevant to epigenetic ageing in common dolphins. This is consistent with the liver functioning as a primary organ for trace element storage and detoxification (Cáceres-Saez et al., 2018; Hansen & Danscher, 1995).

Essential trace elements in humans have a narrow optimal concentration range for physiological function, and both insufficient and excessive levels may adversely affect health (Birgisdottir et al., 2013; Zoroddu et al., 2019). Based on this, one possible explanation for the observed positive associations between EAA and the essential metals zinc and selenium is that concentrations may fall below optimal physiological levels or exceed non-toxic thresholds, both of which could adversely affect biological ageing processes. In contrast, manganese and copper may have occurred within an optimal physiological range, where concentrations are sufficient for normal biological function without inducing toxicity. The decelerated EAA associated with manganese may reflect its role at lower concentrations as a cofactor in antioxidant defence systems (Korc, 1983). This could suggest that some essential metals can exert context-dependent protective effects on biological ageing pathways.

Metal mixture exposure has similarly been associated with increased EAA (Wang et al., 2025) in human studies. More specifically, essential metals may be associated with lower EAA, whereas non-essential metals are more consistently associated with increased EAA (Bozack et al., 2022; Xiao et al., 2021). However, the present findings indicate a more complex pattern, with both positive and negative associations observed among individual essential trace elements. Differences between the present findings and human studies may reflect species-specific variation in metal toxicokinetics and detoxification pathways. Dolphins are exposed to distinct environmental metal mixtures and may exhibit different bioaccumulation patterns and essential metal regulation compared with humans. In addition, differences in exposure levels, and sample size may contribute to the heterogeneous associations observed for essential trace elements across studies.

Importantly, organisms are exposed to complex mixtures of contaminants rather than isolated compounds. For trace elements, interactions among metals can result in antagonistic, synergistic, or additive effects, which differ from the effect of any single metal alone (Braun et al., 2016; Carlin et al., 2013). The mechanisms by which heavy metals affect DNA methylation are not yet understood. Several studies suggest that heavy metals may exert toxic effects via interactions with biomacromolecules (Katti & Igumenova, 2024; Rupa et al., 2023; Warren et al., 1998), induction of oxidative stress and the production of reactive oxygen species (ROS) (Albrecht & Matyja, 1996; DesMarias & Costa, 2019; Hu et al., 2020; Qu & Zheng, 2024) and inhibition of enzymes as well as disruption of metabolic pathways (Fernandes Azevedo et al., 2012; Oluranti et al., 2021; Ran et al., 2022; Samborska et al., 2004; Warren et al., 1998) that could potentially lead to increased EAA.

In our study, mercury measured both in hepatic and renal tissue significantly explained variation in EAA. However, the mechanisms by which mercury influences EAA remain largely unclear. Several studies in humans have shown that mercury exposure can alter DNA methylation patterns at both global and gene-specific levels (Goodrich et al., 2013). As epigenetic clocks are based on DNA methylation, these findings suggest that mercury exposure may plausibly influence EAA by altering DNA methylation levels.

Traditionally, the selenium-to-mercury (Se:Hg) ratio has been used as an indicator of mercury detoxification capacity, with a ratio above one suggesting sufficient selenium availability to facilitate the formation of inert mercuric selenide complexes, thereby reducing mercury toxicity (Law et al., 2003). Conversely, a lower Se:Hg ratio has been interpreted as an indication that mercury exposure may exceed selenium’s protective capacity. However, in our study the Se:Hg ratio was not significantly associated with EAA in either unadjusted or adjusted OLS models, nor in multivariate distance-based linear models (DISTLM). This consistent lack of association suggests that the Se:Hg ratio does not meaningfully explain variation in EAA in common dolphins. One explanation is the Se:Hg ratio may inadequately reflect the complex physiological and molecular processes linking contaminant exposure to biological ageing in common dolphins. This could be particularly pertinent where long-term cumulative exposure, species-specific detoxification mechanisms, and sublethal epigenetic effects are involved.

In addition to trace metal exposure, we assessed the effect of PFAS on EAA. Our analysis of 28 PFAS variables together did not show a significant association with EAA; however, a subset of PFAS variables (PFBA, PFDA, PFHxS-B, and PFNA) did indicate an association with EAA, and remained statistically significant after adjusting for sex, sexual maturity, and the selenium-to-mercury (Se:Hg) ratio. This finding indicates that EAA may be only driven by a small number of PFAS variables. These results differ from a recent study that reported total PFAS burdens were associated with accelerated biological ageing in common dolphins (Peters et al. in review). In that study, the authors showed 1 ng/g increase in total PFAS corresponded to a 0.031-year increase in age acceleration (approximately 11 days), with no effects of sex and the best-fitting model explaining 17.2% of EAA variation (Peters et al. in review). However, finding total versus compound specific differences perhaps is to be expected. Multivariate PFAS models can dilute compound-specific effects. This occurs because many PFAS variables are highly inter-correlated due to shared exposure sources. High variation across several prevalent inter-correlated PFAS variables can obscure the influence of individual compounds within multivariate frameworks. As a result, biologically relevant associations driven by a small number of particularly active compounds may not emerge when all PFAS variables are analysed collectively.

Several factors may explain why our subset of PFAS (PFBA, PFDA, PFHxS-B, and PFNA) showed significant associations with EAA. One likely explanation is that individual PFAS variables differ substantially in their toxicokinetic behaviour, biological persistence, tissue affinity, and capacity to disrupt molecular pathways involved in ageing (Dunmyer et al., 2026; Fischer et al., 2024; Huang et al., 2019). Although PFAS are often grouped collectively, they represent a chemically diverse class of compounds with differing carbon-chain lengths, functional groups, bioaccumulation potential, and biological activities (Buck et al., 2011; Calafat et al., 2007; Eriksson et al., 2017; Gkika et al., 2025). Consequently, only certain compounds may meaningfully influence pathways linked to epigenetic regulation and biological ageing.

For example, longer-chain perfluoroalkyl carboxylic acids such as PFDA and PFNA are known to be particularly bioaccumulative and proteinophilic (Cao et al., 2022; Conder et al., 2008), with strong affinity for liver tissue and relatively long biological half-lives. These compounds have been associated in other taxa with oxidative stress, lipid dysregulation, mitochondrial dysfunction, endocrine disruption, and altered DNA methylation mechanisms that are also implicated in epigenetic ageing processes. Their persistence may, therefore, result in more sustained cellular stress capable of influencing EAA.

In contrast, PFHxS-B (a branched isomer of PFHxS) may reflect differences in isomer-specific toxicodynamics. Branched and linear PFAS isomers can differ in environmental behaviour, protein binding, metabolism, and tissue retention, potentially leading to differing biological effects despite structural similarity (Beesoon & Martin, 2015; Schulz et al., 2020; Wijayahena et al., 2025). Emerging evidence suggests that branched PFAS isomers may exhibit distinct toxicological profiles and differential interactions with molecular signalling pathways (Schulz et al., 2020).

The association observed with PFBA is somewhat more unexpected given its relatively short carbon chain and lower bioaccumulation potential compared with longer-chain PFAS. However, PFBA may act as a marker of more recent or ongoing exposure sources, co-exposure to other unmeasured contaminants, or particular exposure pathways. Specifically, PFBA is associated with a range of industrial and consumer applications, including non-stick and stain-resistant coatings, food contact materials, fire-fighting foams, and various industrial processes. It is also frequently detected as a transformation product resulting from the environmental or metabolic degradation of higher-chain PFAS used in treated textiles, paper-based food packaging, and carpeting materials. Historically, PFBA has additionally been employed in the manufacture of photographic film (Minnesota Department of Health, 2022).

Alternatively, short-chain PFAS may exhibit biological effects disproportionate to their persistence through mechanisms involving cellular signalling, immune modulation, or metabolic disruption. Given the comparatively limited toxicological understanding of short-chain PFAS in marine mammals, this finding warrants further investigation.

In human studies, PFAS exposure has been associated with changes in DNA methylation patterns (Abdulkadir et al., 2025; Schmidt, 2022) and alterations in mitochondrial function (Hofmann et al., 2023; Kam et al., 2025). Furthermore, PFAS exposure has been shown to affect amino acid and lipid metabolism (Guo et al., 2022), disrupt endocrine signalling (Bulawska et al., 2025; Di Nisio et al., 2022) and induce oxidative stress (Iakovou & Kourti, 2022; Solan et al., 2023), which are all pathways linked to biological ageing. Additionally, accelerated biological ageing is strongly linked to chronic inflammation (Li et al., 2023), and there is evidence that PFAS exposure can increase inflammatory responses (Dragon et al., 2023). C-reactive protein (CRP) is a key biomarker involved in chronic inflammation associated with ageing. In addition, both CRP levels and the triglyceride–glucose (TyG) index, a measure combining two lab tests, triglyceride and fasting blood glucose, have been shown to correlate strongly with biological ageing (Flood et al., 2023; He et al., 2024; Lopez-Jaramillo et al., 2023; Yuan et al., 2023). Although the underlying mechanisms of PFAS were not directly assessed in this study, similar inflammation-related pathways may also operate in common dolphins and warrant further investigation.

The best fitting model in our study considered a combined contaminant profile, encompassing the metal subset (Se, Zn, Cu, Al, Mn), the PFAS subset (PFBA, PFDA, PFHxS-B, PFNA), with the Se:Hg ratio, indicating a strong combined effect of contaminants on EAA, with a canonical correlation of 66.7%. This indicates that variation in EAA is better explained by the overall contaminant mixture structure than by individual pollutant classes, highlighting the importance of considering combined exposure profiles in marine mammals.

In the marine environment, trace metals and PFAS are co-exposed and therefore may additively or even synergistically affect common dolphins. A recent study in humans reported that PFAS and heavy metals can interact in complex, dose- and ratio-dependent ways, producing either synergistic or antagonistic toxic effects (Sands et al., 2025). In that study, the authors also found that combined exposure to these contaminants leads to distinct, cell-type–specific responses across different cell lines. Overall, these findings suggest that PFAS–heavy metal mixtures may influence cellular function through toxicological pathways that depend on both the specific composition of the mixture (including dose and ratio) and the type of cell exposed (Sands et al., 2025).

Beyond contaminant influence, sex and sexual maturity showed variable influence on EAA. While there was no significant difference in EAA values, on average, between males and females in our study, the direction of the effect is consistent with studies in humans (Oblak et al., 2021), where males report marginally higher average EAA. One explanation for the higher mean EAA in males compared to females, is the generally shorter lifespans observed in males across many species (Alberts et al., 2014). Another factor potentially explaining why males could exhibit accelerated epigenetic ageing, compared to females, is the maternal offloading of contaminants. Female dolphins can transfer metals and other pollutants to their offspring via placental transfer and lactation, effectively reducing their own contaminant burden over time (Masuda et al., 1978; Wells et al., 2005). In contrast, males accumulate contaminants continuously throughout their lifespan, without such offloading opportunities, which may contribute to higher lifetime exposure and consequently, accelerated epigenetic ageing.

One potential explanation for the higher EAA observed in mature compared to immature individuals is the increased accumulation of metal burdens with age. Consistent with this a study on Commerson’s dolphins (*Cephalorhynchus commersonii*) from the southwestern Atlantic reported higher concentrations of copper (Cu), zinc (Zn), and cadmium (Cd) in sexually mature dolphins, with no sex-related differences observed (Fernandez et al., 2026). In males, sexual maturity is likely associated with increased testosterone production (Sherman et al., 2021), a hormone which has been linked to increased ROS production and reduced antioxidant activity (Agarwal et al., 2005). In females, reproductive processes such as pregnancy and lactation impose substantial energetic and physiological costs, which may similarly increase oxidative load. These maturity-related increases in oxidative stress provide a plausible mechanistic link to accelerated biological ageing, and may contribute to the elevated epigenetic age acceleration (EAA) observed in mature individuals.

### Study considerations

Since metals clearly clustered into two groups by analytical batch, we residualised the trace metal data to account for batch effects. However, such an approach may have also removed some biologically meaningful signals. In addition, our samples were collected between 2000 and 2023, a period during which environmental regulations changed. PFOS-containing firefighting foams were restricted in New Zealand in 2006, and by 2011 all PFOS-containing products were prohibited (Authority Environmental Protection, 2019). By correcting for the analytical batch effect in trace metals, year effects were partially accounted for, as analytical batch and year were strongly correlated. However, year effects were not similarly accounted for in EAA. This may compromise interpretations and inferences.

Because DNA methylation clocks built with different tissue types demonstrate strong differences in predicted biological age (Richardson et al., 2025) and therefore, also different EAA values, the findings of this study should be validated with clocks for common dolphins based on distinct tissue types.

## Conclusion

Our study provides novel evidence that a subset of trace metals (Se, Zn, Cu, Al, and Mn) and PFAS variables (PFBA, PFDA, PFHxS-B, and PFNA), alongside sexual maturity, are associated with variation in EAA in common dolphins. Mature dolphins exhibited significantly higher EAA compared to sexually immature individuals, highlighting the influence of life-history processes on biological ageing. The significant effect of sexual maturity on EAA highlights the importance of incorporating detailed life-history information in future studies. It is possible that these compounds do not directly drive EAA themselves, but instead act as proxies for broader contaminant mixtures, dietary patterns, habitat use, trophic position, or chronic physiological stressors associated with accelerated biological ageing. Importantly, associations between contaminant burdens and EAA remained significant after accounting for sex and sexual maturity, suggesting that exposure to these compounds may contribute to accelerated biological ageing beyond intrinsic factors. Given the complex and poorly understood nature of PFAS toxicology in cetaceans, caution is warranted in attributing causality. Nonetheless, our collective findings support the potential use of EAA as an integrative biomarker linking environmental exposure to the physiology and ageing processes in dolphin populations.

## Acknowledgements

This study was funded by a Royal Society Te Apārangi Rutherford Discovery Fellowship (RM22073) and Massey University Research Funding (RM27575 and RM26712) awarded to KAS. Completion was possible thanks to technical funding support provided by The Colgan Foundation (RM28041). Permits for sample collection (39239-MAR & 111522-MAR) were obtained from the New Zealand Department of Conservation Te Papa Atawhai in partnership with local iwi and hapū (Indigenous New Zealanders). We acknowledge Hannah Hendriks (Department of Conservation Te Papa Atawhai) and Robert Brooke (Epigenetic Clock Development Foundation) who supported the original epigenetics study which underpins the current study. Samples were accessed via BIOCET (The New Zealand Cetacean Biobank) hosted by Massey University. The authors thank members of the Cetacean Ecology Research Group (CERG) and New Zealand Institute of Advanced Studies (NZIAS) for supporting aspects of this work.

## Declaration of generative AI and AI-assisted technologies in the manuscript preparation process

During the preparation of this work, the authors used ChatGPT for assistance to edit and abbreviate text and graphical abstract design, and ChatGPT and Claude for support with coding in R. The authors reviewed and edited the output as needed and take full responsibility for the content of the published article.

## Data availability statement

The R code and underlying dataset used in this study will be openly available https://github.com/XXXXXX on GitHub upon final publication. Our datasets will additionally be released through PRIMER-e as a training resource.

## Author contributions

ACL: Conceptualization, Methodology, Sample collection, Formal analysis, Investigation, Data curation, Visualization, Writing – original draft, Writing – review & editing. EMFH: Conceptualization, Laboratory analyses, Data curation, Writing – review & editing. ELB: Sample collection, Laboratory analyses, Data curation, Writing – review & editing. XS: Laboratory analyses, Data curation, Writing – review & editing. MJA: Formal analysis, Investigation, Visualization, Software, Writing – review & editing. SG: Laboratory analyses, Data curation, Writing – review & editing. SSM: Sample collection, Writing – review & editing. WG: Writing – review & editing. KJP: Writing – review & editing; SY: Writing – review & editing. JJ: Formal analysis, Visualization, Writing – review & editing. KAS: Conceptualization, Resources, Data curation, Laboratory analyses, Funding acquisition, Sample collection, Writing – review & editing.

